# *Lactobacillus reuteri*: direct passage from ingested yogurts to urine microbiota

**DOI:** 10.1101/2019.12.11.872788

**Authors:** Jean-Christophe Lagier, Fatima Mekhalif, Vicky Merhej, Hervé Chaudet, Jérémy Delerce, Anthony Levasseur, Didier Raoult

## Abstract

**Background:** Serendipitously, it was observed that fecal transplantation made for *Clostridium difficile* may cure chronic urinary tract infections. This led us to evaluate the passage in the urine of probiotics contained in yoghurt, which have been claimed to prevent urinary infections.

**Results:** A commercial yogurt that contained 3 probiotics (*Bifidobacterium animalis, Lactobacillus delbrueckiii* and *Lactobacillus reuteri*) was consumed by 28 healthy subjects. We performed by culturomics, urine analysis before and after feeding these yogurts. Genome sequencing of the bacterial strains absent before yogurt consumption and present after consumption was performed. Testing more than 40,000 colonies by MALDI-TOF, we observed in two men and one woman (11% of subjects included), the urine colonization of the same *Lactobacillus reuteri* present in yogurts, with the same genome with 50 genes never identified in other lactobacilli.

**Conclusion:** This confirms that, as ingested, *Lactobacillus salivarius* passes into the milk of lactating women, some bacteria, particularly *Lactobacillus* can colonize body fluids previously considered sterile after ingestion via the digestive route. Although the consequences of this passage remain unknown, we prove for the first time that there is a digestive passage in the urine after consumption of probiotics, including fermented products sold commercially.

## Background

The role of probiotics in the control of urinary tract infections remains controversial [1]. In addition, the mechanism of its putative activity has not yet been clarified. Nevertheless, it was demonstrated that after oral ingestion, *Lactobacillus salivarius* and *L. fermentum* could be identified in human milk [2]; patients treated with these probiotics improved more and had fewer recurrence of mastitis compared to patients treated with antibiotics [2]. In addition, more recently, *L. salivarius* oral administration in pregnant women has also demonstrated its efficiency to prevent infectious mastitis [3]. Therefore, it exists a passage from digestive tract to the biologic fluids. At the same time, a paradigm shift has recently been observed, concomitantly through metagenomics and culture techniques, demonstrating that urine is not sterile [4, 5]. A recent study hypothesizes interconnection from vaginal microbiota and urine bladder microbiota in women by culture. However, it is neglecting oxygen intolerant bacteria which were revealed by metagenomic studies [6] and do not explain male urine microbiota. Here, we have tested the passage of *Lactobacillus* spp. from dairy products after ingestion by healthy individuals. The genome sequences of the cultured strains were analyzed in order to confirm the strain identity.

## Results

In a preliminary work involving 4 different yogurts, we tested after ingestion in urine samples of 8 individuals, 6,144 different colonies identifying 79 different bacterial species (Supplementary table 1). In a 36-year-old man (individual 1), *Lactobacillus reuteri* recovers in the urine sample after 24 hours of Bifidus yogurt consumption (*L. reuteri*, CSUR P4904). Indeed, we selected this yogurt for the second step of our study involving 20 volunteers. Culturomics analysis of this yogurt (Bifidus, Carrefour), identified three different species of probiotics: *Bifidobacterium animalis* (7.10^8^ CFU/mL)*, Lactobacillus delbrueckii* (4.10^7^ CFU/mL) and *Lactobacillus reuteri* (2.10^5^ CFU/mL) (CSUR P4870). We tested the urine of volunteers after 7 days of daily yogurt consumption. We recovered from urine, 34,650 colonies by MALDI-TOF (Table 1). Overall, we identified 151 bacterial species. In 2 individuals, *Lactobacillus reuteri* was absent of the urine sample before yogurts consumption whereas it was present after a week consumption. The 2 individuals were a 30-year-old women (individual 14, *L. reuteri*, CSUR P5461) and a 35-year-old man (individual 25, *L. reuteri*, CSURP5460) (Table 1). Dendrogram performed using the 4 *L.reuteri* strains of this study detected by MALDI-TOF (CSUR4870= yogurt, CSUR 4904= individual 1, CSUR5461= individual 14, CSUR5460=individual 25), as well as 7 other *L.reuteri* strains for which spectra were available in our database revealed 4 different clusters. Three of the strains were included on a same cluster, while the last strain was classified in a close cluster (**Figure 1**).

**Figure 1:**
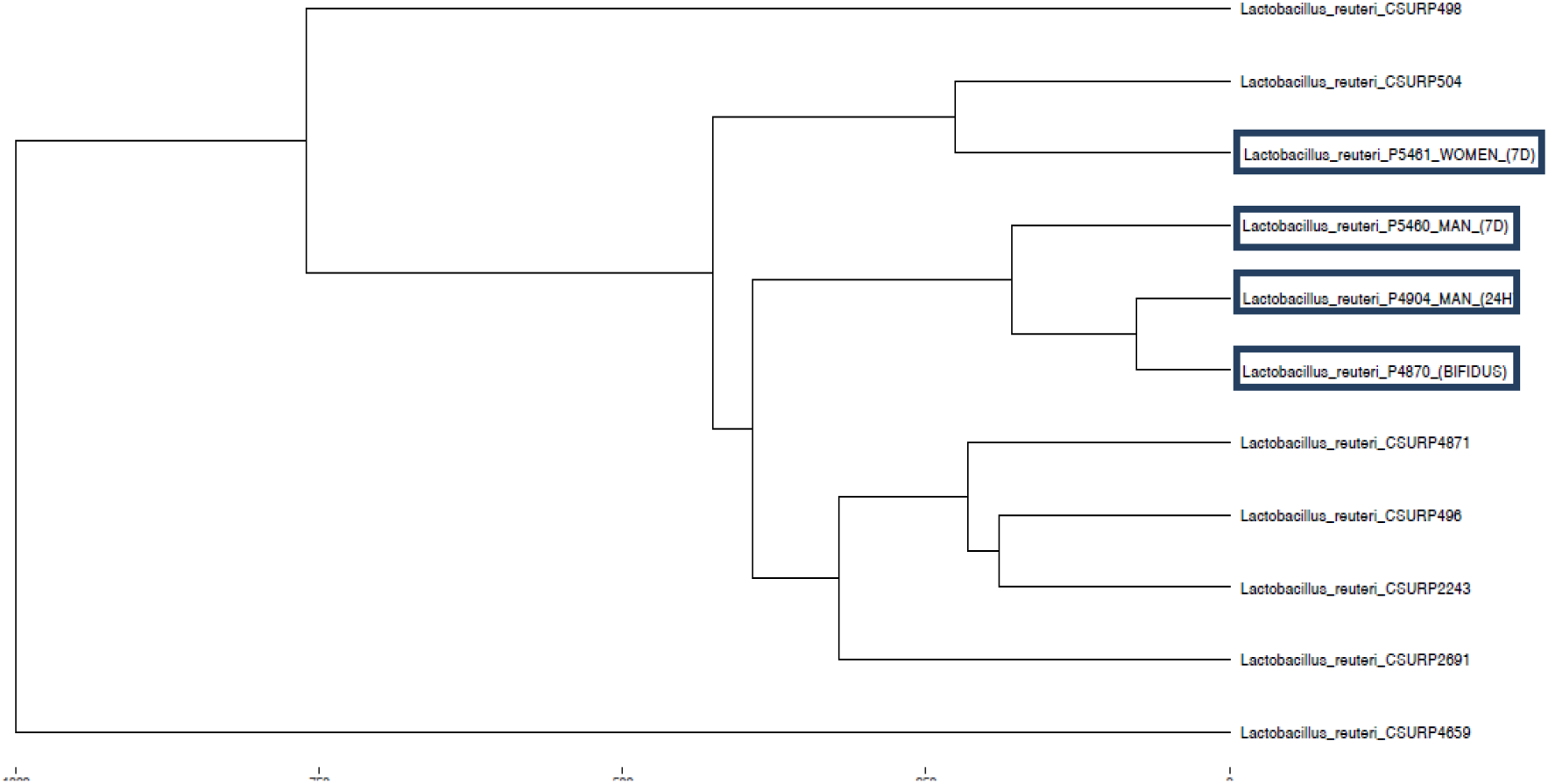
Dendrogram including the 4 *L. reuteri* strains and 7 strains included in our MALDI-TOF database

**Table 1.**
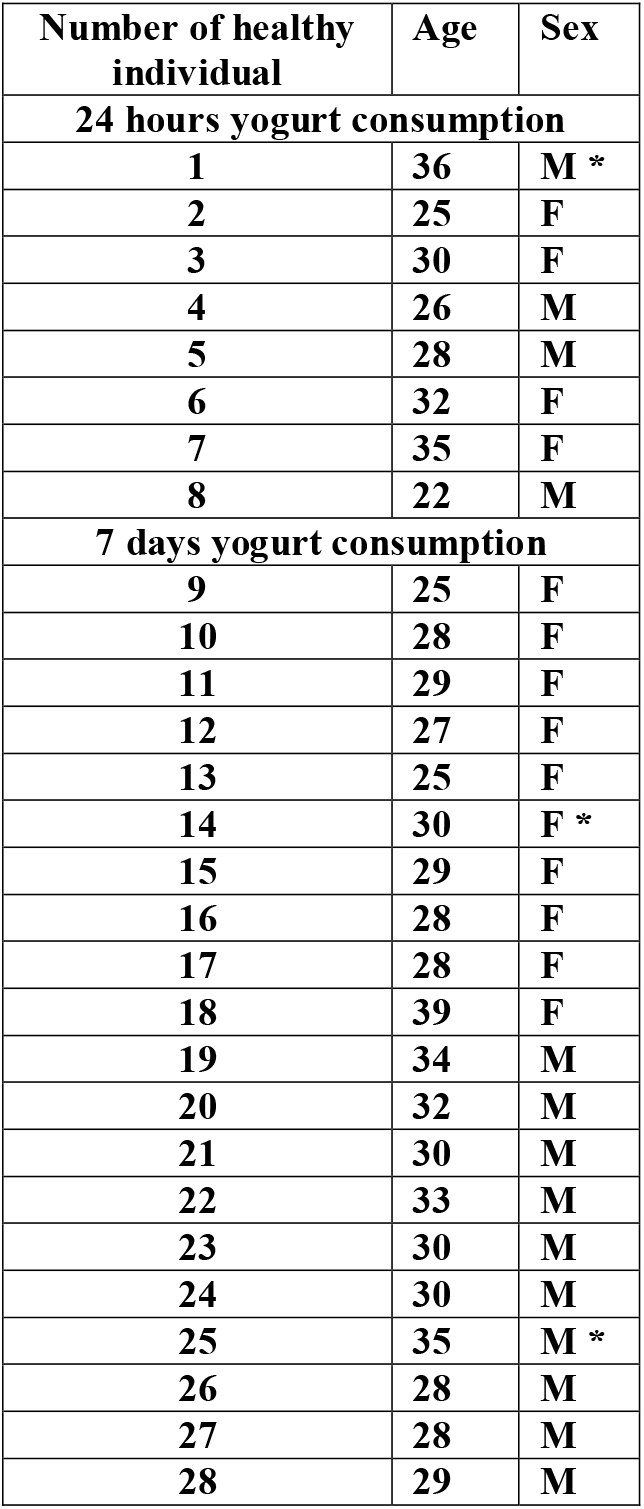

The genomes of the four isolates of *L. reuteri* strains were sequenced and assembled (supplementary table 1). The genome sizes of the four strains were 2,039,275 (+/− 40) bp with a GC content of ~39%. The number of total encoding genes was similar between four strains with an average of 1,631 (+/− 28). In order to study the evolutionary relationships among the different strains of *L. reuteri*, a phylogenetic reconstruction based on the core-genomes alignment of all these strains was carried out (**Figure 2**). The 4 strains of *L reuteri* (CSUR P4870, CSUR P4904, CSUR P5460, CSUR P5461) were distantly separated from all other strains with strongly supported bootstrap values. These 4 strains were grouped along the same branch in a single clade confirming the common origin for these strains. Single nucleotide polymorphisms analyses were performed revealing a close proximity between the strains CSUR P4870, CSUR P4904, CSUR P5460, and CSUR P5461 with only 181 to 432 SNIPS differencing them, as compared to all other strains (**Table 2**). Accordingly, these results confirmed the common origin for these four strains.

**Figure 2.**
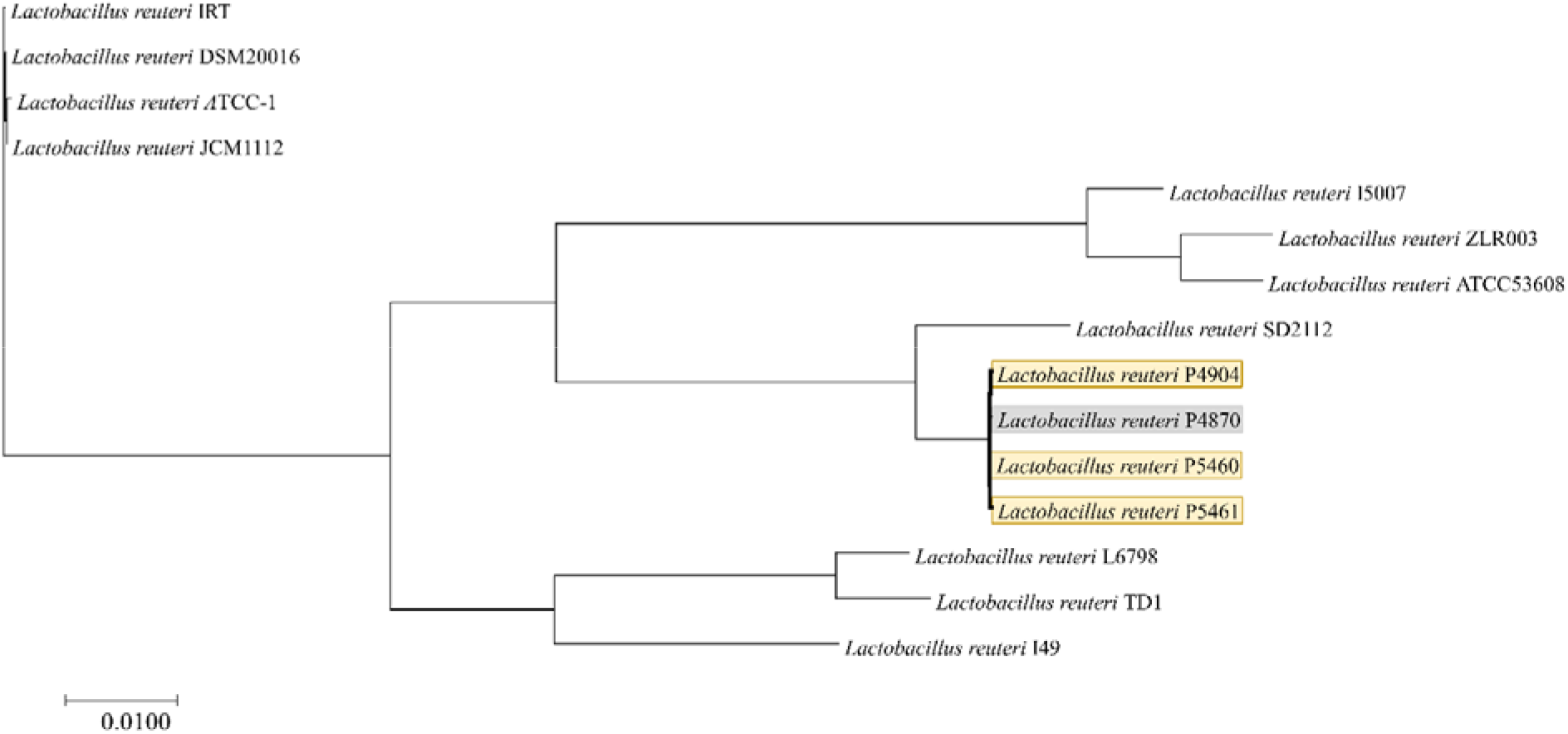
Phylogenetic tree based on core-genome alignment of different *Lactobacillus reuteri* strains.

**Table 2.** Table of SNPs and indels events between *L. reuteri* stains.

|  | *L. reuteri*<br>IRT | *L. reuteri*<br>ATCC-1 | *L. reuteri*<br>DSM20016 | *L. reuteri*<br>JCM1112 | *L. reuteri*<br>ATCC53608 | *L. reuteri*<br>ZLR003 | *L. reuteri*<br>I5007 | *L. reuteri*<br>CSUR<br>P4870 | *L. reuteri*<br>CSUR<br>P4904 | *L. reuteri*<br>CSUR<br>P5460 | *L. reuteri*<br>CSUR<br>P5461 | *L. reuteri*<br>SD2112 | *L. reuteri*<br>I49 | *L. reuteri*<br>L6798 | *L. reuteri*<br>TD1 |
| --- | --- | --- | --- | --- | --- | --- | --- | --- | --- | --- | --- | --- | --- | --- | --- |
| *L. reuteri* IRT |  |  |  |  |  |  |  |  |  |  |  |  |  |  |  |
| *L. reuteri* ATCC-1 | 834 |  |  |  |  |  |  |  |  |  |  |  |  |  |  |
| *L. reuteri* DSM20016 | 475 | 766 |  |  |  |  |  |  |  |  |  |  |  |  |  |
| *L. reuteri* JCM1112 | 619 | 641 | 267 |  |  |  |  |  |  |  |  |  |  |  |  |
| *L. reuteri* ATCC53608 | 60524 | 59331 | 60570 | 60783 |  |  |  |  |  |  |  |  |  |  |  |
| *L. reuteri* ZLR003 | 64523 | 64135 | 63991 | 65175 | 12388 |  |  |  |  |  |  |  |  |  |  |
| *L. reuteri* I5007 | 53033 | 54098 | 54838 | 55358 | 16290 | 15984 |  |  |  |  |  |  |  |  |  |
| *L. reuteri* G1570 | 48731 | 48433 | 47822 | 48957 | 55694 | 55414 | 55561 |  |  |  |  |  |  |  |  |
| *L. reuteri* G1583 | 48825 | 48614 | 47915 | 49044 | 55816 | 55529 | 55665 | 181 |  |  |  |  |  |  |  |
| *L. reuteri* G1614 | 48648 | 48297 | 47703 | 48872 | 55950 | 55332 | 55548 | 432 | 400 |  |  |  |  |  |  |
| *L. reuteri* G1615 | 48727 | 48409 | 47782 | 48952 | 56020 | 55406 | 55610 | 335 | 312 | 206 |  |  |  |  |  |
| *L. reuteri* SD2112 | 66832 | 66796 | 67330 | 67781 | 65786 | 69296 | 68972 | 16364 | 16460 | 16313 | 16361 |  |  |  |  |
| *L. reuteri* I49 | 50930 | 50778 | 50102 | 51303 | 61236 | 64152 | 58355 | 45764 | 45985 | 45815 | 45845 | 58180 |  |  |  |
| *L. reuteri* L6798 | 51812 | 51125 | 51212 | 52295 | 71651 | 64353 | 63457 | 45053 | 45270 | 45154 | 45170 | 57195 | 46452 |  |  |
| *L. reuteri* TD1 | 52276 | 51259 | 51655 | 52743 | 62529 | 72571 | 62734 | 46208 | 46423 | 46367 | 46377 | 59498 | 50280 | 16913 |  |

Orthologous genes were identified by a bidirectional blast approach of all CDS of the 15 genomes of *L. reuteri* using ProteinOrtho (**Figure 3**). This approach detected 1,283 orthologs shared by all 15 genomes, i.e. in average 67% of the CDS of each strain are part of the core proteome. A specific group of 50 encoding-genes was only identified in the *L. reuteri* (CSUR P4870, CSUR P4904, CSUR P5460, CSUR P5461), mostly composed of hypothetical proteins (41 proteins) (**Supplementary Table 1**). Among proteins with assigned functions, we identified proteins involved in metabolism of nitrogen (carbon-nitrogen bonds and other peptide bonds), in transfer of nitrogenous group such as transaminase and in carbohydrate active-transport system. Interestingly, a candidate protein was annotated as an oxalyl-CoA decarboxylase. Finally, all genomes contain two clusters of polyketide synthases and one cluster of nonribosomal peptide synthetases (**Table 3**). The genes order was relatively conserved among genomes and the clusters were found in synthenic regions, with the exception of the genome of *L. reuteri* TD1 that contains one non-modular cluster of polyketide synthases. The lengths of the clusters were very similar among genomes with a mean of 4,148 and 12,566 bp for the polyketide synthases clusters and 3,851 bp for the cluster of nonribosomal peptide synthetases (**Table 3**).

**Figure 3:**
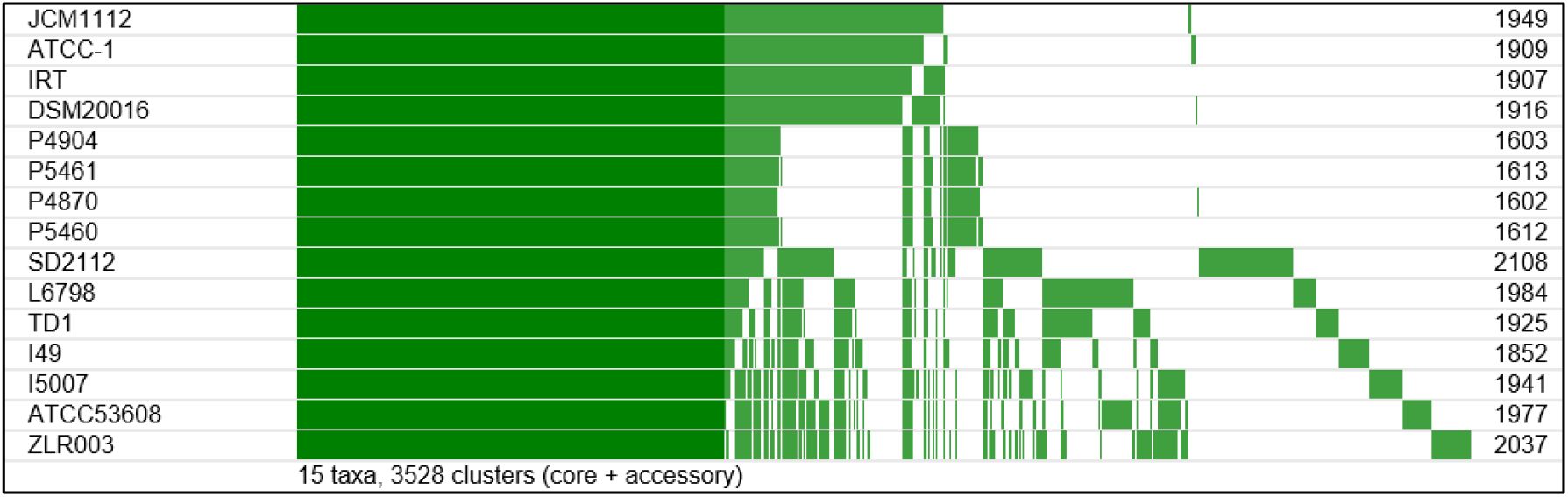
Pan-genome of *L. reuteri* species. Core genome is depicted in dark, accessory genes in light grey. Studied strains of this work are in bold.

**Table 3:** List and number of bacteriocins, polyketide synthases and nonribosomal peptide synthetases detected in the sequenced genomes.

|  | PKS | NRPS | Bacteriocin | name | name | name |
| --- | --- | --- | --- | --- | --- | --- |
| SD2112 | 2 | 1 | 3 | Plantaricin (Lactobacillus plantarum) | circularin_A/uberolysin_family_circular_bacteriocin (Enterococcus faecalis) | Nukacin (Staphylococcus warneri) |
| TD1 | 2 | 1 | 2 | Plantaricin (Lactobacillus plantarum) | circularin_A/uberolysin_family_circular_bacteriocin (Enterococcus faecalis) |  |
| JCM1112 | 2 | 1 | 2 | Plantaricin (Lactobacillus plantarum) | circularin_A/uberolysin_family_circular_bacteriocin (Enterococcus faecalis) |  |
| DSM20016 | 2 | 1 | 2 | Plantaricin (Lactobacillus plantarum) | circularin_A/uberolysin_family_circular_bacteriocin (Enterococcus faecalis) |  |
| G1570 | 2 | 1 | 2 | Plantaricin (Lactobacillus plantarum) | circularin_A/uberolysin_family_circular_bacteriocin (Enterococcus faecalis) |  |
| G1583 | 2 | 1 | 2 | Plantaricin (Lactobacillus plantarum) | circularin_A/uberolysin_family_circular_bacteriocin (Enterococcus faecalis) |  |
| G1614 | 2 | 1 | 2 | Plantaricin (Lactobacillus plantarum) | circularin_A/uberolysin_family_circular_bacteriocin (Enterococcus faecalis) |  |
| G1615 | 2 | 1 | 2 | Plantaricin (Lactobacillus plantarum) | circularin_A/uberolysin_family_circular_bacteriocin (Enterococcus faecalis) |  |

## Discussion

Here, we demonstrated the proof of concept that the consumption of dairy products can lead to the presence of fermenting bacteria in the urine human fluids. We are strongly confident in our data as we included negative and positive controls for each culturomics and genomic step [7]. In addition, we observed the same results in 3 different individuals among the 28 tested (10.7%), including 2 collected in men. This seems to rule out the usual pattern of vaginal contamination of urine with *Lactobacillus*. To the best of our knowledge, none study have previously demonstrated that oral administration of probiotic could be followed by a urinary passage from the gut. Coupled with genome analysis allowing to hypothesize that specific bacterial characteristics facilitated modification of urine microbiota, this opens several reflections on the possible colonization of the internal mucosa by transient bacteria through the digestive tract.

The precise role, the duration of excretion of these *Lactobacillus* in urine after a single dose, and the significance of this passage are not currently fully understandable. The decrease of the diversity of the human urinary microbiota is a risk factor of urinary tract infection. The role of the administration of probiotic in protecting against urinary tract infection in both men and women remains controversial [1, 13]. Nevertheless, intra urethral *Lactobacillus casei* administration improves urinary tract infection caused by *E. coli* in mice [14]. If many questions remain unanswered, our results should help to end the debate over the meaning of *Lactobacillus* in the urine, which has long been thought to be exclusively female and of vaginal origin [6]. The origin from the digestive tract by oral ingestion of yogurts in the urine supposes a direct passage from the digestive tract to the bladder through the blood or lymphatic system and opens a new field of reflection.

Interestingly, oxalyl-CoA decarboxylase is known to be a major enzyme involved in the bacterial detoxification of oxalate within the human intestinal tract [15]. Oxalate is a toxic compound widely distributed in food [16]. The possible role of oxalate-degrading anaerobic bacteria in oxalate-related diseases has already been studied, and a direct correlation between oral administration of probiotic bacteria and the reduction of urinary oxalate excretion has been evidenced in rats and humans [17]. This suggests that administration of probiotics could have a role in the prevention of oxalate urinary lithiasis. *Lactobacillus* spp. have been reported for a long time as having an antimicrobial activity and are acidifying culture media. Both of these actions may be of interest in preventing bladder infections.

## Conclusion

Finally, it was recently demonstrated that fecal microbiota transplantation led not only to the resolution of recurrent *Clostridium difficile* infections, but also to a significantly decrease of recurrent urinary tract infections frequency, and improved antibiotic susceptibility profile of causative microorganisms lacked explanation [18]. The possible passage of microorganisms from digestive tract could lead to a paradigm shift in the understanding of urinary tract infections and beyond that on the relationships between urinary microbiota and human health [1, 19].

## Material and Methods

### Patients and samples

Healthy individuals, aged from 24 to 44 years old, were recruited. Inclusion criteria were absence of urinary tract infection symptoms, absence of treatment with antibiotics in the last month, absence of yogurt consumption in the last 2 days. The first step of the study included 8 individuals (4 male and 4 female). Microbiological analysis of the urine was performed before and after the consumption of 2 yogurts (125g) a day. Two individuals have consumed 4 different yogurts (Bifidus, Carrefour, France; Nature sucré, Carrefour, France, Activia brassé Nature, Danone; Danacol, Danone, France). We selected these yogurts among their large consumption in France and we directly bought in supermarket these yogurts with a date of expiry compatible with their consumption.

The second step of the study included 20 volunteers (10 male and 10 female). Microbiological analysis of the urine was performed before the consumption of one of the selected yogurt (Bifidus, Carrefour, France) and after the consumption of 2 yogurts (125g) by 24 hours during 1 week. Urines were collected after local disinfection. This study was approved by the Ethics Committee of the IHU Méditerranée Infection under Number 2016-011.

### Culturomics analysis

Urine samples, as well as yogurt samples, were directly inoculated in Man Rogosa Sharpe (MRS) agar (Biomérieux, Marcy l’Etoile, France) and in 5% sheep blood agar (Biomérieux, Marcy l’Etoile, France) in both aerobic and anaerobic atmosphere using GENbag anaer system at 37°C. Culture were preserved during 5 days and all the colonies were tested by MALDI-TOF mass spectrometry (Bruker Daltonik, Leipzig, Germany) [7].

### Identification methods

The growing colonies were first identified using our systematic matrix-assisted laser desorption-ionization time-of-flight mass spectrometry (MALDI-TOF-MS) screening on a MicroFlex LT system spectrometer (Bruker Daltonics) [8]. Each deposit was covered with 2 mL of a matrix solution (saturated α-cyano acid-4-hydroxycinnamic in 50% acetonitrile and 2.5% trifluoroacetic acid). For each spectrum, a maximum of 100 peaks was used, and these peaks were compared to those of previous samples in the computer database of the Bruker Base and in our homemade database, including previously identified bacterial spectra. Protein profiles are regularly updated based on the results of clinical diagnoses and on new species, providing new spectra, our database containing 7,462 bacterial spectra at the time of our study.

If, after three attempts, the bacterial colony could not be accurately identified by MALDI-TOF or if the bacterium had not previously been isolated from the human gut, 16S rRNA gene-based identification was performed by amplifying and sequencing the bacterial 16S rRNA gene extracted from the pure culture using the previously described DNA extraction method.

#### Strain deposition, 16S rRNA and genome sequencing accession numbers

All the strains of *Lactobacillus reuteri* isolated in this study were sequenced and deposited in the Collection de Souches de l’Unité des Rickettsies (CSUR, WDCM 875) and are readily available at (http://www.mediterraneeinfection.com/article.php?1aref=14&titre=collection-de-souches&PHPSESSID=cncregk417fl97gheb8k7u7t07) (Supplementary Table 2). The 16S rRNA accession numbers and the draft genomes are deposited with an available GenBank accession number (Supplementary Table 2).

### Genome sequencing

Genomic DNAs of *Lactobacillus reuteri* CSUR*P*4870*, Lactobacillus reuteri* CSUR*P*4904*, Lactobacillus reuteri* CSUR*P*5460 and *Lactobacillus reuteri* CSUR*P*5461 were sequenced on the MiSeq Technology (Illumina, Inc, San Diego CA 92121, USA) with paired end and barcode strategy in order to be mixed with others projects constructed according to the Nextera XT library kit (Illumina). The gDNAs were quantified by a Qubit assay with the high sensitivity kit (Life technologies, Carlsbad, CA, USA) to 12.6 ng/μl, 2.4 ng/μl, 33.9 ng/μl and 23.7 ng/μg respectively, and dilutions were performed to require an input of 1 ng of DNA. The « tagmentation » step fragmented the genomic DNA. Then, limited cycle PCR amplification (12 cycles) completed the tag adapters and introduced dual-index barcodes. After purification on AMPure beads (Lifetechnolgies, Carlsbad, CA, USA), the libraries were then normalized on specific beads according to the Nextera XT protocol (Illumina). Normalized libraries are pooled for sequencing on the MiSeq. Automated cluster generation and paired-end sequencing with dual index reads was performed in a single 39-hour run in a 2×251bp. Within this pooled run, the numbers of paired-end reads were 577,861 for *Lactobacillus reuteri* CSUR*P*4870, 723,210 for *Lactobacillus reuteri* CSUR*P*4904, 1,732,818 for *Lactobacillus reuteri* CSUR*P*5460 and 1,514,107 for *Lactobacillus reuteri* CSUR*P*5461. The paired end reads were trimmed and filtered according to the read qualities.

#### Bioinformatic and evolutionary analyses

All complete genomes from *Lactobacillus reuteri* were downloaded from NCBI: *Lactobacillus reuteri* DSM 20016, GCA_000016825.1; *Lactobacillus reuteri* JCM 1112, GCA_000010005.1; *Lactobacillus reuteri* SD2112, GCA_000159455.2; *Lactobacillus reuteri* ATCC 53608, GCA_000236455.2; *Lactobacillus reuteri* I5007, GCA_000410995.1, *Lactobacillus reuteri* TD1, GCA_000439275.1, *Lactobacillus reuteri* IRT, GCA_001046835.1, *Lactobacillus reuteri* ZLR003, GCA_001618905.1, *Lactobacillus reuteri* I49, GCA_001688685.2. After sequencing, reads from strains *Lactobacillus reuteri* (CSUR4870, CSUR 4904, CSUR5460, and CSUR5461) were mapped on the reference *L. reuteri* JCM 1112 using CLC Genomics workbench 7 with the following thresholds: 0.7 for coverage and 0.8 for similarity. Consensus sequences were extracted from the mapped reads (>3 reads threshold). All genomes were annotated using Prokka using default parameters [9]. For orthologous detection, we applied Proteinortho with default values [10]. All orthologous genes were aligned using Muscle, then concatenated [11]. Phylogenetic reconstruction was performed using RaxML (Randomized Axelerated Maximum Likelihood) with the GTRCAT model and bootstrap value of 100 [12]. Single nucleotide polymorphisms and insertion deletion events were calculated based on aligned orthologous sequences.

### Comparative Genomic and Pan-genome Analysis

Gene prediction and annotation of all genomes were done with Prokka. To estimate the size of the pan-genome among *Lactobacillus reuteri*, their predicted proteins were clustered using the Proteinortho tool [10] using the reciprocal best hits strategy with 1 × 10^−3^, 30% and 70% as thresholds for the BLASTp e-value, and identity and coverage of amino acid sequences, respectively.

---

**Supplementary table 1.** Nucleotide content and gene count levels of the *L. reuteri* genomes (strains CSUR P4870, P4904, P5460 and P5461)

|  | **L. reuteri**<br>**CSUR**<br>**P4870** | **L reuteri**<br>**CSUR**<br>**P4904** | **L reuteri**<br>**CSUR**<br>**P5460** | **L. reuteri**<br>**CSUR**<br>**P5461** |
| --- | --- | --- | --- | --- |
| **Size (bp)** | 2,039,305 | 2,039,310 | 2,039,225 | 2,039,262 |
| **% G-C content** | 39 | 39 | 39,2 | 39,2 |
| **Number total of genes** | 1,679 | 1,676 | 1,740 | 1,740 |
| **Number total of protein encoding genes** | 1,607 | 1,607 | 1,656 | 1,656 |
| **Number total of tmRNA Genes** | 1 | 1 | 1 | 1 |
| **Number total of tRNA Genes** | 65 | 65 | 65 | 65 |
| **Number total of RNA (5S, 16S, 23S) Genes** | 6 | 6 | 18 | 18 |

**Supplementary Table 2:**
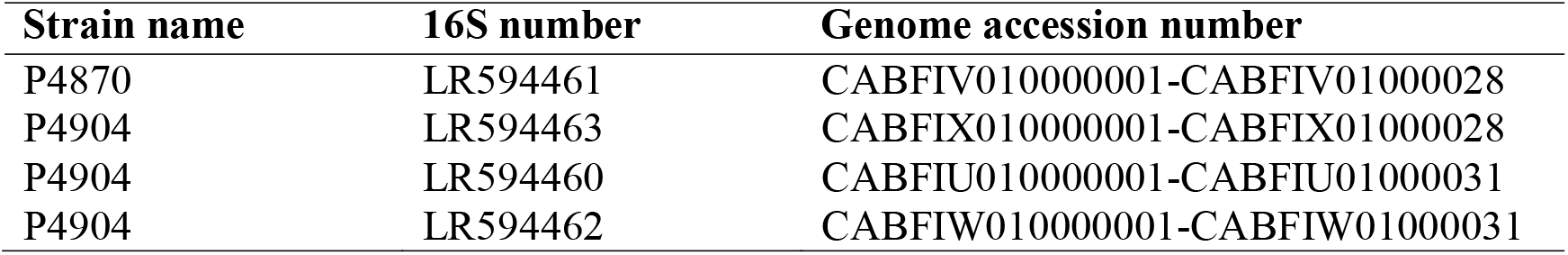
accession numbers of the strains CSUR P4870, P4904, P5460 and P5461

## DECLARATIONS

### Ethical approval and consent to participate

The healthy individuals have consent to participate. This study was approved by the Ethics Committee of the IHU Méditerranée Infection under Number 2016-011.

### Consent for publication

not applicable

### Availability of data and material

Strain deposition numbers, 16S rRNA and genome sequencing accession numbers are available on supplementary Table 2.

### Competing interests

The authors declare that they have no competing interests

### Funding

This work has received financial support from the French Government through the Agence Nationale pour la Recherche, including the “Programme d’Investissement d’Avenir” under the reference Méditerranée Infection 10-IAHU-03.

### Author’s contribution

JCL and DR designed the study; JCL and FM performed the experiments; JCL, FM, VM, HC, JD, AL and DR contributed new reagents/analytic tools; JCL, FM, VM, HC, JD, AL and DR analyzed the data; JCL, FM, AL and DR wrote the paper.

### Competing financial interest

The authors declare no competing financial interests

## Reference List

[1] Whiteside SA, Razvi H, Dave S, Reid G, Burton JP. The microbiome of the urinary tract--a role beyond infection. Nat Rev Urol 2015 Feb; 12(2):81–90.

[2] Arroyo R, Martin V, Maldonado A, Jimenez E, Fernandez L, Rodriguez JM. Treatment of infectious mastitis during lactation: antibiotics versus oral administration of Lactobacilli isolated from breast milk. Clin Infect Dis 2010 Jun 15; 50(12):1551–8.

[3] Fernandez L, Cardenas N, Arroyo R, et al. Prevention of Infectious Mastitis by Oral Administration of Lactobacillus salivarius PS2 During Late Pregnancy. Clin Infect Dis 2016 Mar 1; 62(5):568–73.

[4] Hilt EE, McKinley K, Pearce MM, et al. Urine is not sterile: use of enhanced urine culture techniques to detect resident bacterial flora in the adult female bladder. J Clin Microbiol 2014 Mar; 52(3):871–6.

[5] Wolfe AJ, Toh E, Shibata N, et al. Evidence of uncultivated bacteria in the adult female bladder. J Clin Microbiol 2012 Apr; 50(4): 1376–83.

[6] Thomas-White K, Forster SC, Kumar N, et al. Culturing of female bladder bacteria reveals an interconnected urogenital microbiota. Nat Commun 2018 Apr 19; 9(1):1557.

[7] Lagier JC, Khelaifia S, Tidjani-Alou M. Culture of previously uncultured members of the human gut microbiota by culturomics. Nat Microbiol. [in press] 2016.

[8] Reid G, Bruce AW, Fraser N, Heinemann C, Owen J, Henning B. Oral probiotics can resolve urogenital infections. FEMS Immunol Med Microbiol 2001 Feb; 30(1):49–52.

[9] Asahara T, Nomoto K, Watanuki M, Yokokura T. Antimicrobial activity of intraurethrally administered probiotic Lactobacillus casei in a murine model of Escherichia coli urinary tract infection. Antimicrob Agents Chemother 2001 Jun; 45(6):1751–60.

[10] Lung HY, Cornelius JG, Peck AB. Cloning and expression of the oxalyl-CoA decarboxylase gene from the bacterium, Oxalobacter formigenes: prospects for gene therapy to control Caoxalate kidney stone formation. Am J Kidney Dis 1991 Apr; 17(4):381–5.

[11] Holmes RP, Goodman HO, Assimos DG. Contribution of dietary oxalate to urinary oxalate excretion. Kidney Int 2001 Jan; 59(1):270–6.

[12] Turroni S, Vitali B, Bendazzoli C, et al. Oxalate consumption by lactobacilli: evaluation of oxalyl-CoA decarboxylase and formyl-CoA transferase activity in Lactobacillus acidophilus. J Appl Microbiol 2007 Nov; 103(5):1600–9.

[13] Tariq R, Pardi DS, Tosh PK, Walker RC, Razonable RR, Khanna S. Fecal Microbiota Transplantation for Recurrent Clostridium difficile Infection Reduces Recurrent Urinary Tract Infection Frequency. Clin Infect Dis 2017 Oct 30; 65(10):1745–7.

[14] Raoult D. Is there a link between urinary microbiota and bladder cancer? Eur J Epidemiol 2017 Mar; 32(3):255.

[15] Seng P, Abat C, Rolain JM, et al. Identification of rare pathogenic bacteria in a clinical microbiology laboratory: impact of matrix-assisted laser desorption ionization-time of flight mass spectrometry. J Clin Microbiol 2013 Jul; 51(7):2182–94.

[16] Seemann T. Prokka: rapid prokaryotic genome annotation. Bioinformatics 2014 Jul 15; 30(14):2068–9.

[17] Lechner M, Findeiss S, Steiner L, Marz M, Stadler PF, Prohaska SJ. Proteinortho: detection of (co-)orthologs in large-scale analysis. BMC Bioinformatics 2011 Apr 28; 12:124.

[18] Edgar RC. MUSCLE: multiple sequence alignment with high accuracy and high throughput. Nucleic Acids Res 2004; 32(5):1792–7.

[19] Stamatakis A. RAxML version 8: a tool for phylogenetic analysis and post-analysis of large phylogenies. Bioinformatics 2014 May 1; 30(9):1312–3.

